# A Pore-Forming Toxin Monalysin Contributes to Infection-Induced Suppression of Defecation in Female *Drosophila*

**DOI:** 10.64898/2025.12.03.692093

**Authors:** Marko Rubinić, Kathirvel Alagesan, Igor Iatsenko

## Abstract

Pathogen expulsion from the gut via defecation is an important defence strategy against infection. The microbial factors that can subvert this defence reaction remain poorly understood. While many microbes have been found to increase host intestinal peristalsis, *Pseudomonas entomophila* infection in *Drosophila melanogaster* leads to infection-induced defecation blockage, particularly in females. Here, we show that this phenotype is driven by a secreted, thermosensitive protein regulated by the GacS/GacA two-component system. Proteomic comparison of the Δ*gacA* mutant, which does not inhibit defecation, and the avirulent Δ*hfq* mutant lacking RNA chaperon Hfq, which still triggers the phenotype, identified pore-forming toxin Monalysin as one of the candidate factors required for inhibiting defecation. Consistently, the Monalysin-deficient mutant was unable to inhibit defecation. Hence, Monalysin besides causing intestinal damage, has a previously unknown role in suppressing defecation and potentially pathogen expulsion. Overall, our study identified a bacterial factor that rapidly reduces defecation frequency, consistent with transient suppression of intestinal transit, thus advancing our understanding of pathogen strategies used to subvert host defences.

## INTRODUCTION

Intestinal peristalsis is a series of wave-like muscle contractions that move food and other gut content through the gastrointestinal tract. The primary role of intestinal peristalsis is to facilitate food digestion and nutrient absorption ^1^. Waste removal and the eviction of toxins, parasites, and pathogens are additional equally important functions of intestinal peristalsis. Intestinal motility is controlled by neuronal regulation and hormonal signaling and can be affected by environmental conditions such as diet and microbial exposure ^2–5^.

Intestinal motility plays an important role in host-microbiota interactions and host defense against infections ^6,7^. For instance, some commensal bacteria, such as *Lacticaseibacillus rhamnosus* ^8^ and *Lacticaseibacillus casei* ^9^ have been found to increase intestinal motility in mammals. Such an effect of commensal bacteria on host intestinal peristalsis is mediated either by a direct response to microbial metabolites and cell wall components, or indirectly through intestinal immune activation by microbial associated molecular patterns ^7^. In mice, microbiota has been shown to provide the host with resistance against intestinal helminths via its ability to increase intestinal motility ^10^. These and other studies identified a number of microbial molecules, including neurotransmitter acetylcholine and toxins driving diarrhea, that increase intestinal peristalsis ^11^. However, much less is known about the microbial determinants that suppress the host intestinal peristalsis.

*Drosophila melanogaster,* thanks to its extensive genetic toolbox and manipulatable microbiota, has been used to study the molecular mechanisms of host-microbiota interactions ^12–15^. Given that intestinal motility in *Drosophila* operates via principles similar to mammals, *Drosophila* provides the intestinal model system to study the general mechanisms underlying the effect of microbes on the host intestinal peristalsis ^16,17^. Several studies explored the effect of certain pathogens and commensals on *Drosophila* gut motility. For example, the Gram-negative bacterium *Pectobacterium carotovorum* –*Ecc15* secretes uracil in the gut ^18^. The presence of uracil triggers a host immune reaction leading to the DUOX-dependent production of reactive oxygen species (hypochlorous acid, HOCl) which is released in the lumen ^18^. Du and colleagues have shown that the evolutionarily conserved Transient Receptor Potential A1 channel (TRPA1) binds HOCl and triggers increased defecation ^17^. Benguettat et al ^19^ extended these findings and proposed that the binding of HOCl to TRPA1 induces a calcium flux in anterior Diuretic Hormone 31 (DH31)-positive enteroendocrine cells, thus promoting DH31 release. DH31 in turn binds to its receptor, DH31-R, in the neighboring visceral muscle and triggers contractions. Such visceral muscle contractions help the host to evict opportunistic bacteria from the gut via defecation, demonstrating that pathogen expulsion via defecation is a defense mechanism against intestinal pathogens ^17,19^. The same mechanism likely explains the increased defecation rate of flies infected with *Pseudomonas aeruginosa* or colonized with specific microbiota members, like *Enterococcus haemoperoxidus* ^16^. Another *Drosophila* commensal *Lactiplantibacillus plantarum* was shown to promote gut motility in larvae via production of acetylcholine – a neurotransmitter with a known role in enabling muscle movement ^20^. While several bacteria have been reported to increase intestinal peristalsis and defecation in flies, whether and how pathogens suppress this defense reaction has not been elucidated.

Recently, we showed that *Pseudomonas entomophila*, a natural pathogen of *Drosophila* ^21^, inhibits defecation and prevents pathogen clearance from intestine in female but not in male flies ^22^. Given that the enforcement of defecation frequency with chemical or genetic tools protects flies from infection ^22^ indicates that such blockage of defecation is an essential previously unknown aspect of *P. entomophila* pathogenesis. Several of *P. entomophila* virulence factors have been identified, including the pore-forming toxin Monalysin and secreted protease, AprA, which allows *P. entomophila* to counteract the intestinal immune response ^23,24^*. P. entomophila* virulence is controlled by two global regulatory systems: the GacS/GacA two component system ^21,25^, and a second system called *Pseudomonas virulence factor* (*pvf*) – a gene cluster which encodes a secreted secondary metabolite ^26,27^. However, *P. entomophila* factors inhibiting defecation and pathogen eviction remain unknown. Identification of these factors will advance our understanding of pathogen strategies used to subvert host defences.

In the present study, we used biochemical, genetic, and proteomic approaches to identify pathogen factors contributing to infection-induced suppression of defecation and pathogen clearance. We showed that the responsible component is a secreted, thermosensitive protein regulated by the GacS/GacA two component system. We employed proteomic approach to compare proteins produced by different *P. entomophila* strains that differently alter defecation in female flies. Finally, we identified Monalysin as one of the *P. entomophila* virulence factors required to suppress pathogen eviction via defecation.

## RESULTS

### A secreted, proteinaceous factor from *P. entomophila* is sufficient to reduce defecation

To identify *P. entomophila* factor(s) causing defecation blockage, we utilized a defecation assay previously used by us and others ^17,22^. We fed flies with bacteria or culture supernatant mixed with a blue color dye that allowed us to estimate the defecation frequency by scoring the number of blue-colored defecation spots deposited within the scoring period of 2 h (Fig. 1A). Using this assay, we first tested whether *P. entomophila* must be alive to block defecation. As shown in Fig. 1B, defecation frequency of flies fed with heat-inactivated *P. entomophila* was significantly higher as compared to flies fed with alive pathogen, although still lower than in control flies (Fig. 1B, Fig. 1B’). These results suggest that at least some of the *P. entomophila* factors involved in defecation blockage are heat sensitive. Next, we investigated the effect of *P. entomophila* cell-free culture supernatant on defecation to test whether a secreted factor is sufficient to reduce defecation. Indeed, we found that flies fed with cell-free supernatant exhibited the same degree of reduction in defecation as flies fed with the pathogen (Fig. 1C, Fig. 1C’). Heat treatment of the cell-free supernatant abolished its ability to inhibit defecation (Fig. 1D, Fig. 1D’). Finally, incubation with Proteinase K had a similar effect (Fig. 1E, Fig. 1E’), supporting a proteinaceous effector. Together, these results identify the active component as a secreted thermosensitive proteinaceous factor that triggers infection-induced defecation blockage.

**Figure 1:**
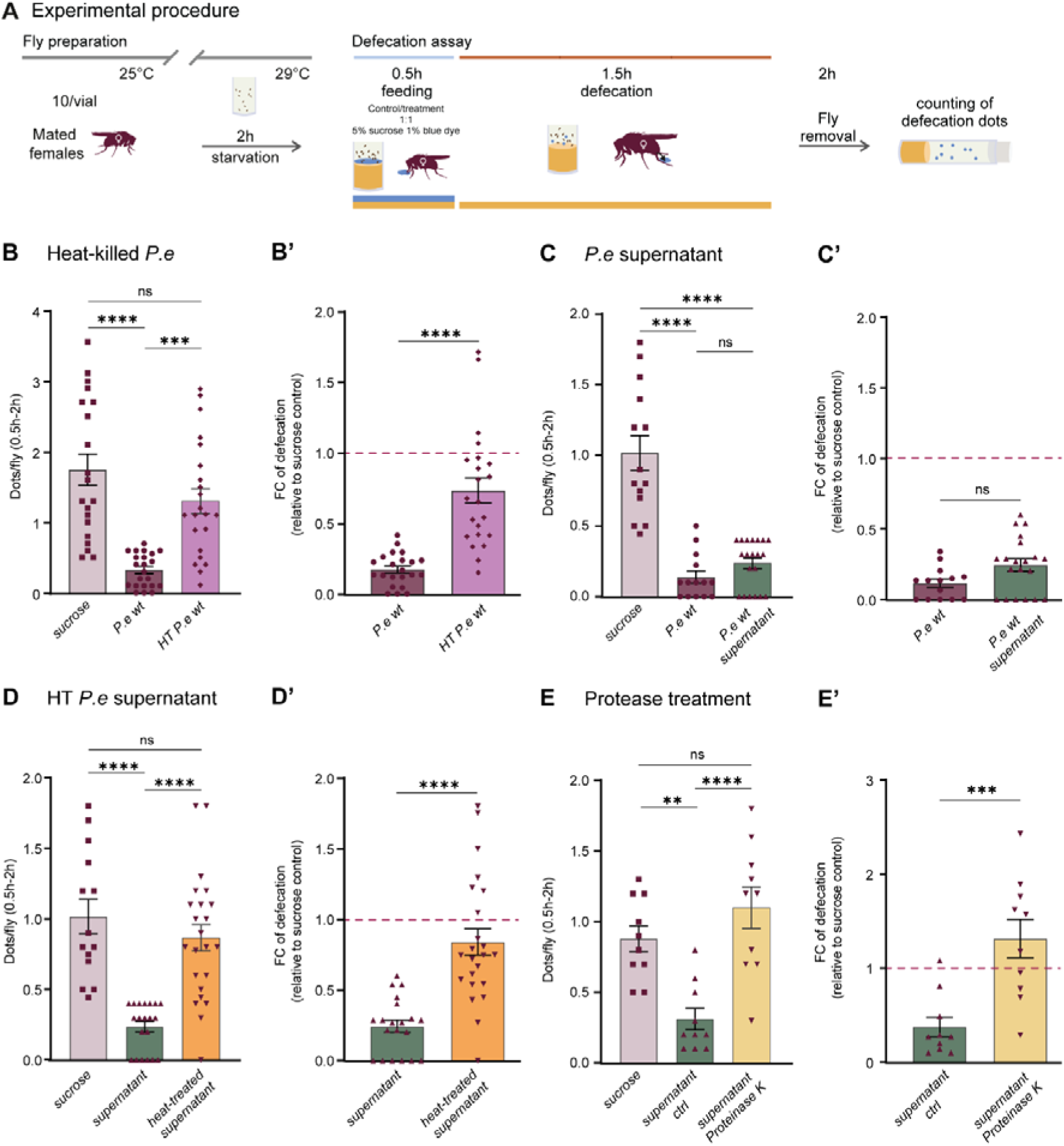
A secreted *P. entomophila* protein is sufficient to reduce defecation. **(A)** Graphical illustration of defecation assay. **(B and B’)** The defecation rate of female *w1118* iso flies measured 0.5 to 2 h after exposure to blue-dyed PBS (sucrose control), *P. entomophila* or heat-killed *P. entomophila* (HT) in 0.5 h protocol. **(B)** Significance by one-way ANOVA with Šídák’s multiple comparisons test. (n (sucrose) = 20, n (*P.e* wt) = 22, n (HT *P.e* wt) = 22, N (days) = 4). **(B’)** Fold change calculated from mean values of PBS control of appropriate treatment from the same biological repeat. Significance by Mann Whitney test. **(C and C’)** The defecation rate of female *w1118* iso flies measured 0.5 to 2 h after exposure to blue-dyed LB (sucrose control), *P. entomophila* or *P. entomophila* cell-free culture supernatant in 0.5 h protocol. **(C)** Significance by one-way ANOVA with Šídák’s multiple comparisons test. (n (sucrose) = 14, n (*P.e* wt) = 14, n (*P.e* wt supernatant) = 20, N (days) = 4). **(C’)** Fold change calculated from mean values of LB control of appropriate treatment from the same biological repeat. Significance by Mann Whitney test. **(D and D’)** The defecation rate of female *w1118* iso flies measured 0.5 to 2 h after exposure to blue-dyed LB (sucrose control), *P. entomophila* cell-free culture supernatant, or heat-treated *P. entomophila* cell-free culture supernatant in 0.5 h protocol. **(D)** Significance by one-way ANOVA with Šídák’s multiple comparisons test. (n (sucrose) = 14, n (supernatant) = 20, n (heat-treated supernatant) = 23, N (days) = 4). (D’) Fold change calculated from mean values of LB control of appropriate treatment from the same biological repeat. Significance by Mann Whitney test. **(E and E’)** The defecation rate of female *w1118* iso flies measured 0.5 to 2 h after exposure to blue-dyed LB (sucrose control), *P. entomophila* cell-free culture supernatant, or *P. entomophila* cell-free culture supernatant treated with Proteinase K in 0.5 h protocol. **(E)** Significance by one-way ANOVA with Šídák’s multiple comparisons test. (n (sucrose) = 10, n (supernatant ctrl) = 10, n (supernatant Proteinase K) = 10, N (days) = 3). **(E’)** Fold change calculated from mean values of LB control of appropriate treatment from the same biological repeat. Significance by Mann Whitney test.

### The GacS/GacA system controls secretion of the defecation-blocking factor

We next examined bacterial regulatory systems that could control the secretion of the active peptide/protein. We started with the RNA chaperone Hfq which plays important regulatory role in bacteria ^28^. Given that we previously showed that Hfq responds to the gut environment and is required for *P. entomophila* virulence ^22^, we tested whether Hfq is necessary for inhibition of defecation. As shown in Fig 2A and 2A’, infection with Δ*hfq* mutant inhibited defecation to the same degree as infection with wild-type *P. entomophila.* Hence, while Hfq is necessary for virulence, it is not required for defecation blockage. Next, we tested GacS/GasA system known to control secondary metabolite production, protein secretion, and virulence. The Δ*gacA* mutant failed to suppress defecation (Fig 2B, Fig. 2B’), indicating that the GacA regulatory system governs secretion or production of a protein factor required for infection-induced defecation blockage in female *Drosophila*.

**Figure 2:**
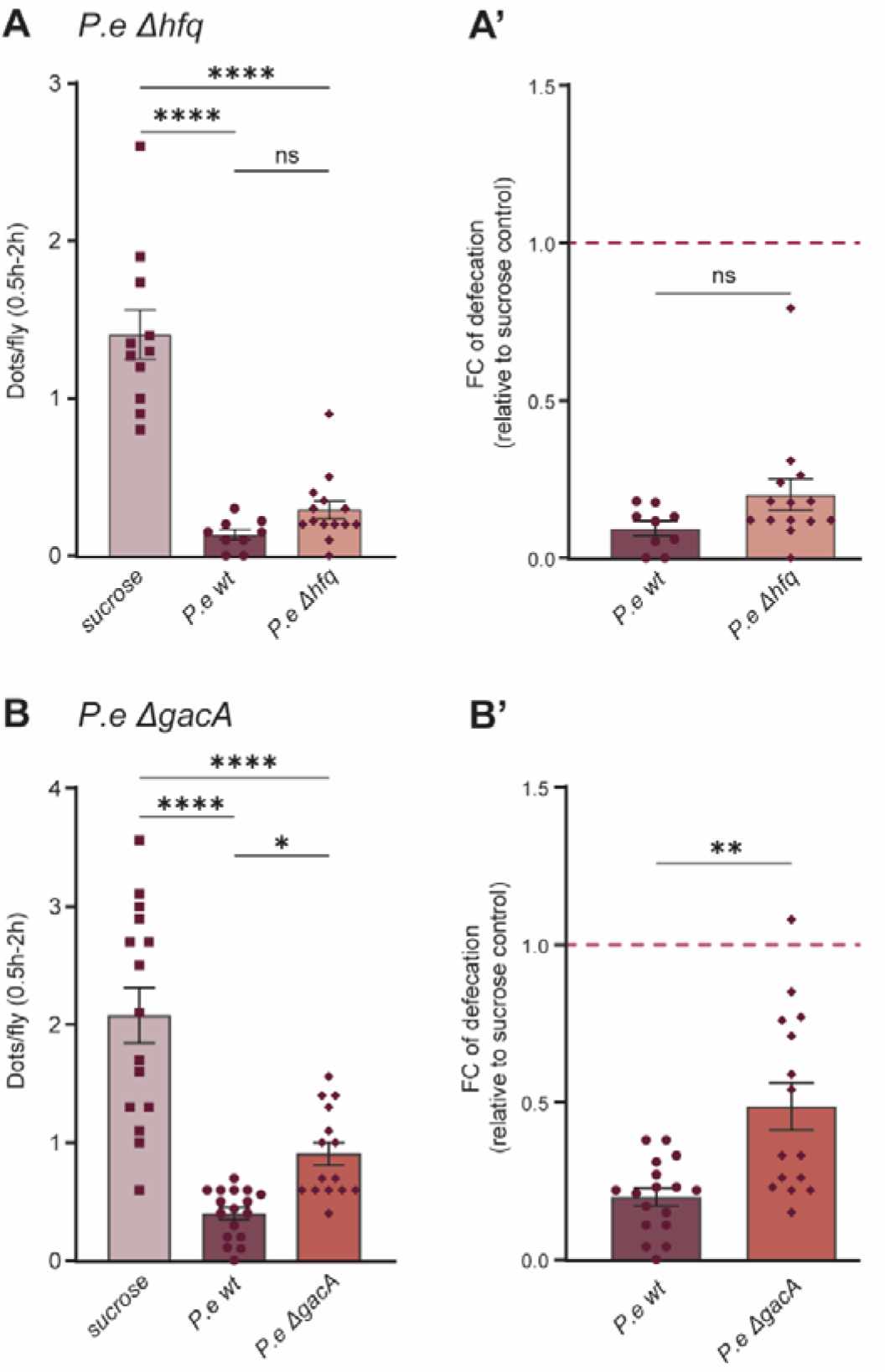
The GacS/GacA system controls secretion of the defecation-blocking factor. (A and A’) The defecation rate of female *w1118* iso flies measured 0.5 to 2 h after exposure to blue-dyed PBS (control), *P. entomophila* wt, or *P. entomophila* Δ*hfq* in 0.5 h protocol. **(A)** Significance by one-way ANOVA with Šídák’s multiple comparisons test. (n (sucrose) = 11, n (*P.e* wt) = 19, n (*P.e* Δ*hfq*) = 14, N (days) = 3). **(A’)** Fold change calculated from mean values of PBS control of appropriate treatment from the same biological repeat. Significance by Mann Whitney test. **(B and B’)** The defecation rate of female *w1118* iso flies measured 0.5 to 2 h after exposure to blue-dyed PBS (control), *P. entomophila* wt, or *P. entomophila* Δ*gacA* in 0.5 h protocol. **(B)** Significance by one-way ANOVA with Šídák’s multiple comparisons test. (n (sucrose) = 15, n (*P.e* wt) = 17, n (*P.e* Δ*hfq*) = 15, N (days) = 4). **(B’)** Fold change calculated from mean values of PBS control of appropriate treatment from the same biological repeat. Significance by Mann Whitney test.

### Comparative proteomics identified Hfq- and GacA-regulated proteins

To proceed further with the identification of protein(s) causing defecation suppression, we compared proteomes of *P. entomophila* wild-type, Δ*gacA,* and Δ*hfq* mutants grown in LB medium. Comparing these three strains allows us 1) to identify GacA- and Hfq-controlled proteins, 2) decouple virulence and defecation-inhibiting factors, as while both Δ*gacA* and Δ*hfq* mutants are avirulent, only Δ*gacA* is impaired in the ability to inhibit defecation.

Using a log2 fold-change cutoff of 1.5, we identified 531 proteins with increased and 108 with decreased abundance in the Δ*hfq* mutant relative to wild-type *P. entomophila* (Fig. 3A, Sup Table 1). We noticed several virulence factors, such as protease AprA, an esterase EstA, flagellin FliC, Pseudomonas virulence factor PvfA, PvfB, PvfC among proteins with increased abundance in Δ*hfq* mutant. To obtain an overview of the functionality of the proteins up-regulated in Δ*hfq* mutant, we performed gene ontology (GO) annotation enrichment analysis. The following GO terms were identified (Fig. S1A): Localization and transport (incl. transmembrane and ABC transporter activity), organic acid and small molecule metabolism, oxidoreductase activity, catabolic processes (organic substance, organonitrogen compound, carboxylic acid), fatty acid metabolism and secondary metabolite biosynthesis. Hence, loss of Hfq increases abundance of proteins linked to transport and metabolic turnover. Several virulence factors, such as catalase KatA ^29^, pilin pilH ^30^, isocitrate lyase AceA ^31^, alginate biosynthesis proteins AlgA, AlgD, AlgE, AlgZ ^32^ were detected among proteins down-regulated in Δ*hfq* mutant (Fig. 3A, Sup Table 1). The following GO terms were enriched among the down-regulated proteins (Fig. S1B): cell envelope organization, oxidoreductase activity acting on metal ions, response to stress, starch and sucrose metabolism, ferroxidase activity. Thus, reduced abundance of proteins involved in stress response and redox balance, suggests that Hfq contributes to oxidative and cell envelope stress response.

**Figure 3:**
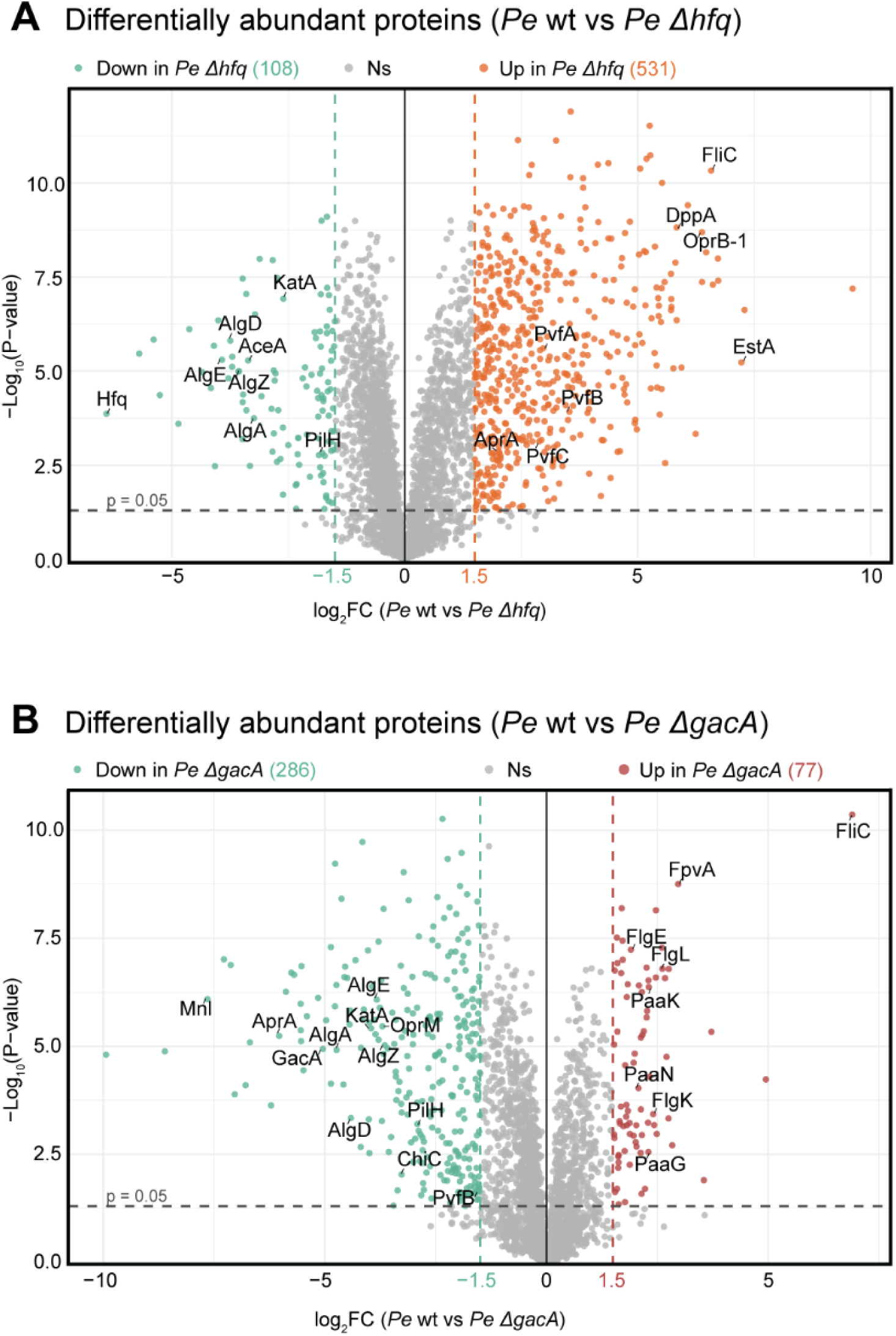
Comparative proteomics identified Hfq- and GacA-regulated proteins. (A) Volcano plots of differentially abundant *P. entomophila* proteins (│log2FC│≥ 1.5 and padj cut-off 0.05) between *P. entomophila* Δ*hfq* and *P. entomophila* wt. (n = 5 independent samples). (B) Volcano plots of differentially abundant *P. entomophila* proteins (│log2FC│≥ 1.5 and padj cut-off 0.05) between *P. entomophila* Δ*gacA* and *P. entomophila* wt. (n = 5 independent samples).

In Δ*gacA* mutant, we found 77 proteins with increased and 286 proteins with decreased abundance as compared to wild-type *P. entomophila* (Fig. 3B, Sup Table 1). Several flagella proteins (FliC, FlgL, FlgK, FlgE) ^33^ and ferric pyoverdine receptor FpvA ^34^ were among virulence-related factors upregulated in Δ*gacA* mutant. Enrichment analysis identified GO terms related to locomotion (Fig. S1C): taxis and flagellar assembly, locomotion and bacterial chemotaxis, chemotaxis signaling, suggesting that deficiency in GacA increases abundance of proteins with a role in motility and chemotaxis. Loss of GacA resulted in a downregulation of a number of virulence factors, including protease AprA, toxin Monalysin, alginate biosynthesis proteins AlgA, AlgD, AlgE, AlgZ, catalase KatA, pilin pilH, chitinase ChiC, Pseudomonas virulence factor PvfB (Fig. 3B, Sup Table 1). GO analysis identified an enrichment of proteins related to secretion, carbohydrate metabolism, and biofilm formation among the down-regulated proteins (Fig. S1D), suggesting that GacA positively regulates secretion systems and metabolic pathways, promoting bacterial adaptation and stability.

Overall, our proteomic analysis not only determined proteins controlled by the two major regulatory systems, GacA and Hfq, but also identified GacA-controlled candidate factors that may inhibit defecation.

### Monalysin is one of the factors inhibiting defecation

To narrow the list of candidates involved in defecation suppression, we sorted the proteins based on the following assumptions. First, the involved protein should be absent/downregulated in Δ*gacA* mutant, which does not cause defecation blockage. Therefore, we performed an additional comparison of the Δ*hfq* mutant with the Δ*gacA* mutant to look for additional proteins that are not present in the mutant that does not cause defecation blockage (Fig. S2A). Second, the involved protein should be present in both *P. entomophila* wild-type and Δ*hfq* mutant, which inhibit defecation. Therefore, we examined the overlap of (1) the protein list that showed no significant difference between *P. entomophila* wild-type and *P. entomophila* Δ*hfq* (indicating that it is present at similar levels in both strains, Fig. 3A) and the protein list that was present in either (2) *P. entomophila* wild-type or (3) *P. entomophila* Δ*hfq* but not in *P. entomophila* Δ*gacA*. Using this approach, we identified 146 candidate proteins that meet our criteria (Fig. 4A). Most of the proteins are uncharacterized with predicted or hypothetical functions. However, we noticed Monalysin as one of the most downregulated proteins in Δ*gacA* mutant (Fig. 4B). It caught our attention, as it is a pore-forming toxin previously shown to be involved in *P. entomophila* pathogenesis^23^. Given that pore-forming toxins are known to interfere with gut peristalsis ^35^, we decided to test the contribution of Monalysin to defecation suppression by *P. entomophila*. Although Δ*mnl* mutant was still able to reduce defecation frequency in infected flies as compared to uninfected controls, this reduction was not as strong as in flies infected with wild-type *P. entomophila* (Fig. 4C, Fig. 4C’). Hence, Monalysin is one of the GacA-regulated proteins that partially mediates defecation blockage caused by *P. entomophila*.

**Figure 4:**
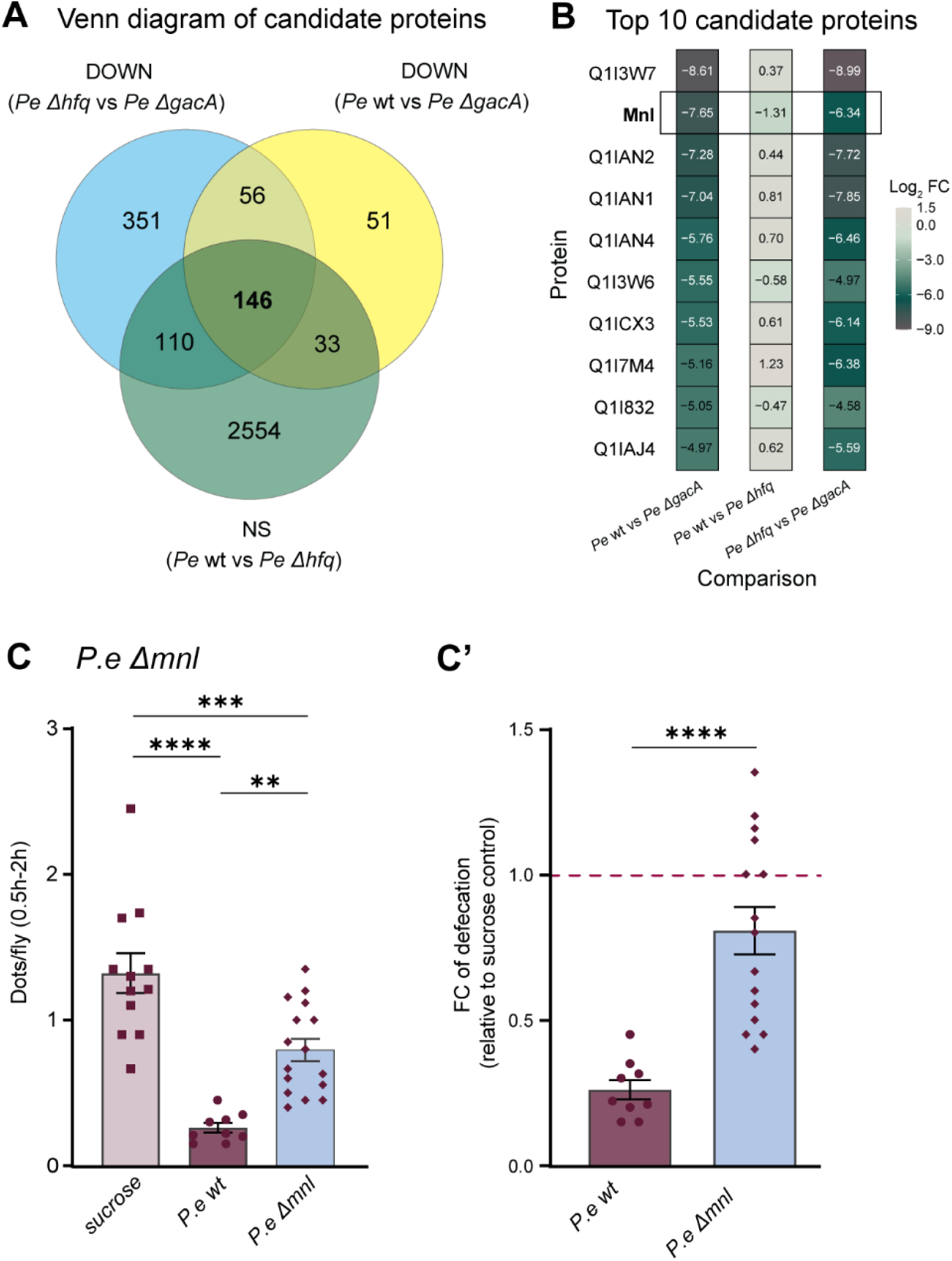
Monalysin is one of the factors inhibiting defecation. (A) Venn diagram showing the overlap of candidate proteins identified in three datasets: Down (*P.e* wt vs *P.e* Δ*gacA*), Down (*P.e* Δ*hfq* vs *P.e* Δ*gacA*), and Ns (*P.e* wt vs *P.e* Δ*hfq*). **(B)** Heatmap showing abundance difference (log2FC) of top 10 candidate proteins (based on *P.e* wt vs *P.e* Δ*gacA* log_2_FC values) in three different comparisons: *P.e* wt vs *P.e* Δ*gacA*, *P.e* Δ*hfq* vs *P.e* Δ*gacA*Δ*gacA*, and *P.e* wt vs *P.e* Δ*hfq*. Significance (padj < 0.05, |log2FC| > 1.5) indicated by *. The highlightied protein is Monalysin. **(C and C’)** The defecation rate of female *w1118* iso flies measured 0.5 to 2 h after exposure to blue-dyed PBS (control), *P. entomophila* wt or *P. entomophila* Δ*mnl* in 0.5 h protocol. **(C)** Significance by one-way ANOVA with Šídák’s multiple comparisons test. (n (sucrose) = 12, n (*P.e* wt) = 9, n (*P.e* Δ*mnl*) = 16, N (days) = 3) **(C’)** Fold change values were calculated as ratios of the corresponding treatment relative to the mean PBS control from the same biological repeat. Significance by Mann Whitney test.

## DISCUSSION

Our study identified a previously unrecognized bacterial factor contributing to infection-induced suppression of defecation frequency in *Drosophila melanogaster*. While several microbes have been shown to accelerate gut motility, *P. entomophila* induces the opposite effect — a marked reduction in defecation, particularly in females. We show that this phenotype does not depend solely on bacterial viability but rather on a secreted, thermosensitive protein whose production is regulated by the GacS/GasA system, and not by the RNA chaperone Hfq.

By combining defecation assays and proteomic analyses, we identified Monalysin as one of the bacterial factors required for suppressing defecation and bacterial clearance. Monalysin is a pore-forming toxin previously shown to be responsible for damage of host intestinal cells^23^. Monalysin also contributes to translation blockage ^36^ and thinning of the intestinal epithelium associated with *P. entomophila* infection ^37^. Our findings expand the role of Monalysin in *P. entomophila* pathogenicity by demonstrating that Monalysin also perturbs gut homeostasis in a way that reduces intestinal transit. The mechanism by which Monalysin causes rapid inhibition of defecation remains to be explored. It could be that intestinal damage inflicted by Monalysin is sufficient to stop peristalsis, as has been suggested for Cry toxins ^38,39^. However, the fact that Δ*mnl* mutant was still able to reduce defecation frequency, although not as strongly as wild-type bacteria, suggests that additional factors besides Monalysin are required to block intestinal peristalsis. Our proteomic comparisons uncovered additional GacA-dependent proteins that may contribute to the phenotype. These factors could act synergistically to modify host gut physiology, either by amplifying epithelial stress or modulating host signaling pathways. Our work, by identifying these proteins provides a starting point for future studies aiming to dissect the bacterial strategies that manipulate host intestinal peristalsis.

Previously, *P. entomophila* was shown to cause food intake cessation in *Drosophila* larvae^24^. Our results provide indirect evidence that such an effect might be caused by Monalysin via inhibition of intestinal peristalsis. Another *Drosophila* pathogen, *Ecc15*, requires *evf* (*Erwinia* virulence factor) gene for bacterial persistence in the larval gut. Although the mechanism of action of *evf* is not known, *evf* was proposed to inhibit gut peristaltic movements and, hence, facilitate bacterial persistence ^40,41^. Pore-forming Cry toxins produced by insecticidal *Bacillus thuringiensis* strains have also been implicated in the feeding cessation and suppression of peristalsis ^38,39^. Hence, abrogation of peristalsis is frequently observed in insects during intestinal infections. On one hand, inhibition of peristalsis would allow the host to reduce the amount of ingested pathogen, repair the midgut epithelium, and reduce the transmission of a pathogen via fecal route. Additionally, trapping the microbes in a specific gut region can help the host to kill the pathogens by antimicrobial peptides ^42^. On the other hand, inhibition of peristalsis would prevent pathogen eviction from intestine via defecation, facilitating bacterial persistence ^22,42^. The facts that pathogens have developed strategies to abrogate peristalsis and hosts with defects in intestinal transit suffer more from infections ^35,43^, suggest that pathogens benefit from inhibiting gut transit. However, our results show that virulence and inhibition of peristalsis are not necessarily linked. The Δ*hfq* mutant, despite being avirulent, still inhibits defecation. This indicates that defecation blockage is not directly associated with virulence or host killing but may represent an adaptive bacterial strategy. Reduced gut motility could prolong bacterial presence in the intestine, enhance colonization, or alter the gut environment in ways favorable for the pathogen. Our findings also open broader questions about how non-pathogenic bacteria affect intestinal peristalsis and whether inhibition of defecation is inherently detrimental to the host. In some cases, reduced motility might limit bacterial dissemination or moderate immune activation, while in others it could promote bacterial retention and chronic infection. The presence of this phenotype in avirulent strains suggests that modulation of gut transit may be a widespread microbial strategy rather than a purely pathogenic one.

Altogether, our findings describe a bacterial factor that suppresses the host’s intestinal peristalsis and the ability to evict bacteria from the gut via defecation. Further elucidation of the underlying mechanisms will not only broaden our understanding of pathogen strategies used to subvert host defences but could also open the way to new methods of insect pest control.

## METHODS

### Arrive protocol

The ARRIVE 2.0 guidelines were followed in planning, conducting and reporting *in vivo* experiments ^44^.

### Drosophila husbandry

*Drosophila* stocks were raised in a light:dark cycle (12 h:12 h) at 25°C, on a standard cornmeal/agar medium (6.2 g agar, 58.8 g cornmeal, 58.8 g inactivated dried yeast, 26.5 ml of a 10% solution of methyl-paraben in 85% ethanol, 60 ml fruit juice, 4.8 ml 99% propionic acid for 1 L). For maintaining flies, flies were transferred to fresh vials every 2-3 days, and fly density was kept to a maximum of 15 flies per vial. At day 2 after eclosion, all emerged adults were transferred to new vials where they were left to mate for additional 2 days. Female flies were then sorted using CO_2_ in vials with 10 or 20 flies per vial the evening before experiment. Therefore, 5-7 days old mated female *w1118* iso flies were used in all experiments.

### Bacterial strains

*Pseudomonas entomophila* L48 strain ^21^, *P. entomophila* Δ*gacA* mutant ^24^, and *P. entomophila* Δ*mnl* ^23^ were obtained from Bruno Lemaitre’s lab. Previously published *P. entomophila* Δ*hfq* mutant ^45^ was kindly provided by Dr. Edna Bode (Max Planck Institute for Terrestrial Microbiology).

### Preparation and treatments of bacteria and bacterial supernatant

*P. entomophila* strains utilized in this study were grown directly from frozen 50% glycerol stocks. Briefly, 10 µL of the stocks was incubated in 20 mL of LB medium in a 50 mL flask overnight (∼16 h, 29°C, 175 rpm). The following day, the overnight culture was diluted 1:16 in 150 mL of LB medium and incubated in 500 mL flasks under the same conditions for a minimum of an additional 24 hours. Before the defecation experiment, 45 mL of bacterial culture was centrifuged at 3,500 × g for 15 min at 4 °C using 50 mL screw-top tubes (114 × 28 mm, PP; Sarstedt AG & Co.).

a. For preparing bacterial suspensions, the supernatant was discarded and the bacterial pellets were resuspended in PBS. The final concentration was adjusted to an optical density (OD600) of 200 in PBS.
b. For preparing cell-free supernatant, after centrifugation of bacterial culture, 2 mL of the supernatant (taken from the top) was filtered using a Millex filter unit (0.22 µm, sterile; Merck Millipore).
c. For heating treatments, 500 µL of bacterial suspension or supernatant was placed in a 1.5 mL Eppendorf vial, heated for 15 min at 95°C (Eppendorf ThermoMixer), and then left to cool for 15 min at room temperature before next steps.
d. For proteinase K treatment, the supernatant was mixed with either control (same volume of ddH_₂_O) or proteinase K (AM2548, Ambion, final concentration: 1 mg/mL) and incubated for 30 min at 37 °C (Eppendorf ThermoMixer) ), and then left to cool for 15 min at room temperature before next steps.

### Quantifying defecation in adult flies

For quantifying defecation, the bacterial suspension or supernatant (described above) were mixed in a 1:1 ratio with a 5% sucrose solution containing 1% blue dye (Food Blue No. 1, TCI) (referred to as blue mixture) as described previously ^17^. Prior to the defecation experiment, 150 µL of blue mixture was pipetted into vials containing standard food and filter papers (referred to as infection vials). Before transferring mated flies to the infection vials, the flies were kept in empty vials for 2 hours at 29°C (dry starvation). Flies were kept in the infection vials for 0.5 h before being transferred to the vial containing conventional non-colored food for an additional 1.5 hours. Defecation quantity was measured by counting ‘defecation dots’ left dried on the inner wall of vials and normalized per number of flies in vial. All experiments were performed at 29°C and at least three times (different days), with a total of at least 9 independent vials per condition, each containing 10 or 20 flies.

### *P. entomophila* Proteome Comparison

Using 1.5 mL Eppendorf tubes, 1 mL of bacterial culture (OD600 = 1, adjusted in LB) was centrifuged at 3,500 × g for 15 min at 4 °C, after which 950 µL of the supernatant was discarded. The pellet was resuspended, and 50 µL of the suspension was transferred to tubes containing glass beads and 500 µL of protein extraction buffer (100 mM Tris-Cl, pH 8.0, 2% SDS, protease inhibitor (cOmplete, Roche, 1 tablet per 4 mL of buffer)) on ice. Following incubation (95°C for 2 minutes) and homogenization (6000 rpm for 30 seconds, Precellys homogenizer), samples were centrifuged (max speed, 10 min, 4°C). Supernatant was collected into Eppendorf tubes. Protein concentrations were measured by the Pierce BCA Protein Assay Kit (Thermo Fisher Scientific Pierce™ BCA Protein Assay Kit Catalog Numbers 23225 and 23227) according to the manufacturer’s protocol. Briefly, 10 μL of samples were incubated with 200 μL of working reagent and incubated for 30 min at 37°C. Absorbance measurements (in duplicates) were performed at 562 nm using a plate reader Infinite M Plex Microplate Reader (Tecan). Samples were stored at -80°C before being analyzed.

#### (a) Sample preparation (SP3)

All samples were subjected to SP3 sample preparation protocol ^46^. A total of 10 μg protein from each sample was mixed with 4× lysis buffer (50 mM HEPES [pH 8; VWR], 4% SDS (vol/vol) (Applichem), 160 mM 2-chloroacetamide (CAA), 40 mM Bond-Breaker Tris(2-carboxyethyl)phosphine hydrochloride [TCEP] solution [Thermo Fisher Scientific]) to a final concentration of 1× absolute volume and incubated for 5 min at 95°C. Samples were cooled to room temperature, and Benzonase nuclease (0.5 units per μg of protein, Merck) was added to each sample. Nucleic acids were degraded during incubation for 30 min at 37°C while shaking at 500 rpm. SP3 beads (1:1 mixture of hydrophobic and hydrophilic carboxyl-coated Sera-Mag SpeedBeads catalog no. 45152105050250 and 65152105050250, GE Healthcare) were added to the samples in a bead/protein ratio of 10:1 (wt/wt). Samples were mixed by pipetting, and anhydrous acetonitrile (Thermo Fisher Scientific) was added to a final concentration of 70% (vol/vol). To allow protein aggregation on the beads, the samples were incubated for 20 min at room temperature. Subsequently, the beads were collected on a magnetic stand for 5 min and the supernatants were discarded. The beads were washed 3 times with 80% (vol/vol) ethanol for 3 min. Beads were air-dried at room temperature for 10 min to allow residual ethanol to evaporate. The beads were thereafter resuspended in 30 μl digestion buffer (50 mM triethylammonium bicarbonate) containing trypsin (Serva) and LysC (Wako) in a 1:50 (wt/wt) enzyme-to-protein ratio. Protein digestion was carried out at 37°C for 14 h in a PCR cycler. The beads were collected on a magnetic stand for 5 min, and the supernatants containing the peptides were collected. The peptide supernatant was then acidified with 2% ACN and 0.1% trifluoroacetic acid.

Label-free DIA analyses of peptides were acquired over 120 min by an Orbitrap Exploris 480 (Thermo Scientific) coupled to a 3000 RSLC nano UPLC (Thermo Scientific) from 500Lng of peptides. Samples were loaded on a PepMap trap cartridge (300 µm i.d.L×L5Lmm, C18, Thermo Scientific) with 2% acetonitrile, 0.1% TFA at a flow rate of 20LµL/min. Peptides were separated over a 25 cm analytical column (PepSep C18, 75Lµm I.D., 1.5 µm). Solvent A consists of 0.1% formic acid in water. Elution was carried out at a constant flow rate of 250LnL/min within 120 min. Initially, a two-step linear gradient was applied: 3–30% solvent B (0.1% formic acid in 80% acetonitrile) within 70.5 min, 30–45% solvent B within 13 min, followed by column washing and equilibration. The column was kept at a constant temperature of 50 °C. The MS was operated in DIA mode for single-injection quantitative measurements of individual samples with the following settings: 60k MS1 resolution, MS1 scan range 350–1250Lm/z, 15k MS2 resolution, MS2 scan range 110–1600Lm/z, Normalised AGC target of 1000%, maximum injection time 50 ms, and fixed normalised collision energy of 30. 10 m/z precursor isolation windows with optimized window placements from 390.4273 to 1210.8002 m/z.

Raw data analysis was performed using Spectronaut (Biognosys AG, Zurich, Switzerland) version 20.2.250922.92449 in directDIA+ deep mode with reviewed UniProt databases (Pseudomonas entomophila (strain L48) - Proteome ID UP000000658, 5,126 entries). Methionine oxidation and Acetyl (Protein N-term) were set as a variable, and carbamidomethylation on cysteine residues was used as a static modification. The FDR for PSM-, peptide-, and protein-level was set to 0.01. All tolerances were set to dynamic for pulsar searches. We characterized differences in proteomic profiles of different bacterial strains. The differential analysis of quantitative proteomics data was performed in Perseus v2.0.7.0. Significance cut-offs were padj < 0.05, |log2FC| > 1.5. Overlap of data was done using VENNY 2.1 (https://bioinfogp.cnb.csic.es/tools/venny/index.html). The R packages ggplot2 was used for data visualization.

GO enrichment analysis was performed by copying the lists of significantly upregulated or downregulated proteins from each dataset into ShinyGO v0.85.1 ^47^. The species was specified as *Pseudomonas entomophila* (taxonomy ID: 384676) using the STRING-db annotation (STRING v12.0). The background list consisted of all proteins detected in the corresponding comparison. Analyses were performed using default settings (FDR cutoff of 0.05, 20 pathways to display, and the option to remove redundancy.

The mass spectrometry proteomics data have been deposited to the ProteomeXchange Consortium via the PRIDE partner repository with the dataset identifier: (*will be included before publication)*.

### Quantification and statistical analysis

Statistical parameters and tests are shown in the figure legends. No formal randomization method was used, but to reduce the potential bias, control and treatment groups were processed in parallel, and sample sizes were balanced. Blinding was not implemented in this study. No data were excluded from analysis. Data analyses and visualization of defecation data were performed using GraphPad Prism 10 software. Proteomics visualization was performed with the R packages ggplot2, dplyr, and tidyverse. Statistical significance was determined using either the Mann Whitney test orone-way, followed by post-hoc tests for multiple comparisons (indicated in the figure legends). Significance was set at p < 0.05, with data presented as mean ± SEM. Significance: *p < 0.05, **p < 0.01, ***p < 0.001, ****p < 0.0001.

## Acknowledgments

We thank Edna Bode and Helge Bode for kindly providing *P. entomophila* Δ*hfq* mutant. We thank Florian Kondrot (MPUSP) for providing technical support for the mass spectrometry experiments. We thank Diane Schad for help with preparation of Graphical Experimental Procedure.This work was supported by the Max Planck Society. I.I. also acknowledges the funding from the Deutsche Forschungsgemeinschaft (grants IA 81/2-1 and IA81/3-1) and from the Boehringer Ingelheim Foundation.

## Author Contributions

I.I. initiated the study and acquired funding. M.R. and I.I. designed the experiments. M.R. performed the experiments. M.R., K.A., and I.I. analyzed the data. I.I. supervised M.R. M.R. and I.I. wrote the manuscript with input from K.A.

---

**Supplementary Figure 1:**
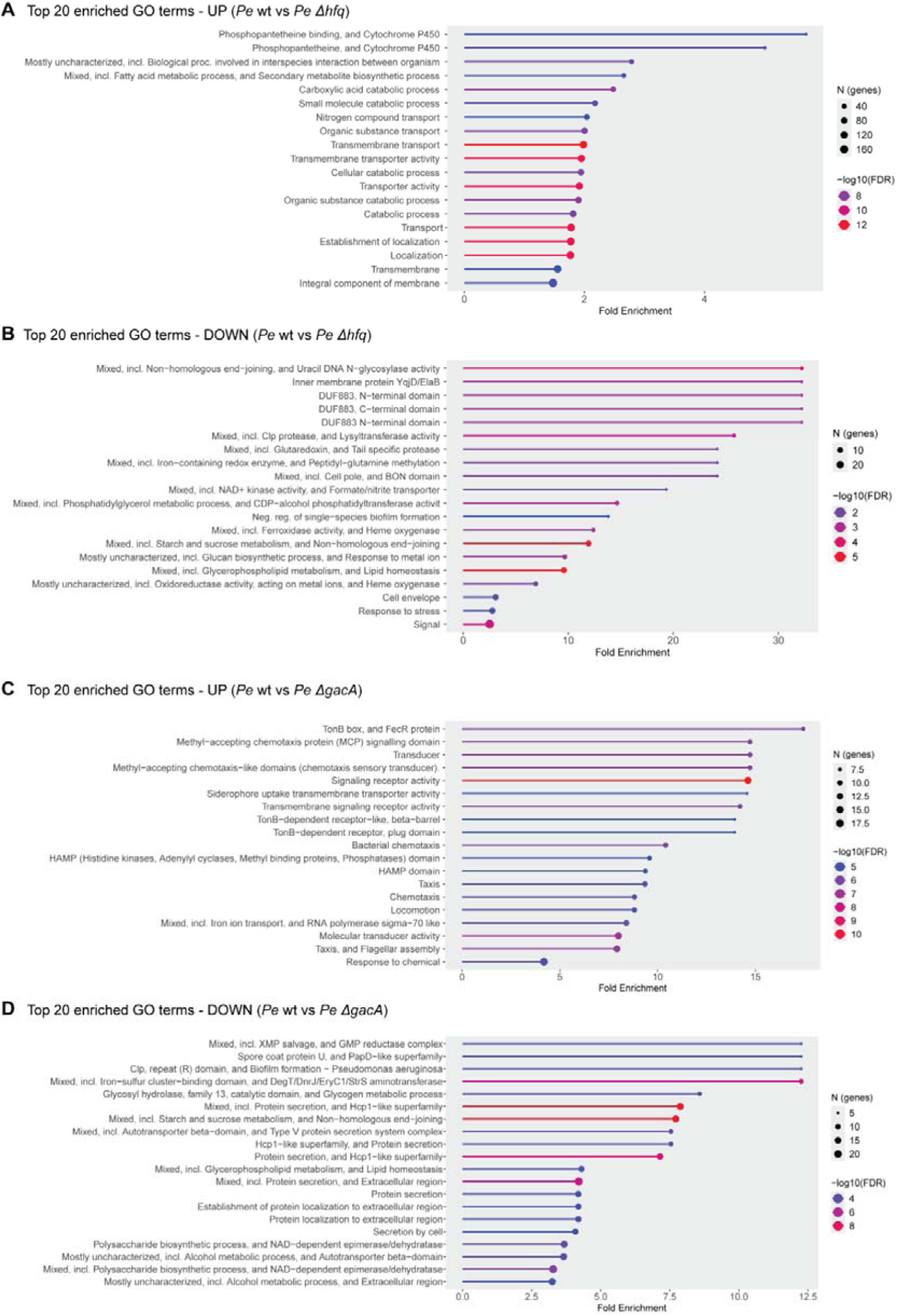
Gene Ontology (GO) enrichment analysis of Hfq- and GacA-regulated proteins in *P. entomophila*. **(A and B)** GO enrichment analysis of proteins differentially abundant in *P. entomophila* Δ*hfq* relative to *P. entomophila* wt. Panels show the top enriched GO terms (p < 0.05) are shown for **(A)** upregulated proteins and **(B)** downregulated proteins identified in the *P. entomophila* Δ*hfq*. **(C and D)** GO enrichment analysis of proteins differentially abundant in *P. entomophila* Δ*gacA* relative to *P. entomophila* wt. Panels show the top enriched GO terms (p < 0.05) are shown for **(C)** upregulated proteins and **(D)** downregulated proteins identified in the *P. entomophila* Δ*gacA*.

**Supplementary Figure 2:**
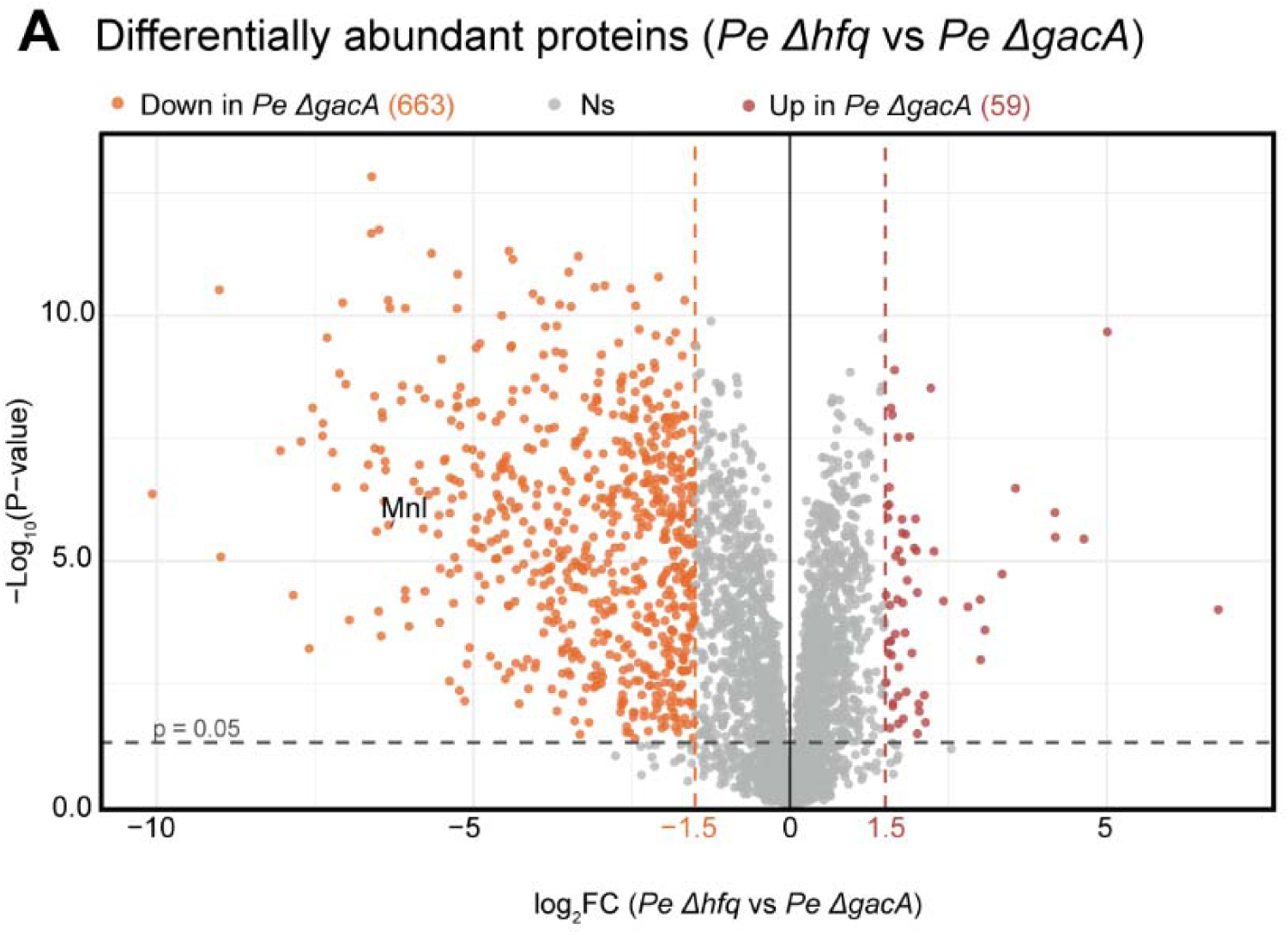
Differences between *P. entomophila* Δ*hfq* and *P. entomophila* Δ*gacA*Δ*gacA* proteomic profiles. **(A)** Volcano plots of differentially abundant *P. entomophila* proteins (│log2FC│≥ 1.5 and padj cut-off 0.05) between *P. entomophila* Δ*hfq* and *P. entomophila* Δ*gacA*. (n = 5 independent samples).

**Supplemental Information**

Table S1. Excel tables listing *P. entomophila* proteins with differential abundance between strains.

